# The role of hnRNP A3 on age-related increased expression of FIX in the Liver

**DOI:** 10.1101/2023.04.22.536972

**Authors:** Toshiyuki Hamada, Hiroko Kushige, Shiori Fukushima, Sumiko Kurachi

**Affiliations:** Department of Pharmaceutical Sciences, International University of Health and Welfare, Ohtawara, Tochigi, 324-8501, Japan; Department of Biological Response and Regulation, Faculty of Health Sciences, Hokkaido University, Sapporo, Hokkaido, 060-0812, Japan; Hakujikai Institute of Gerontology, 5-11-1, Shikahama, Adachi Ward, Tokyo, 123-0864, Japan; Age Dimension Research Center, National Institute of Advanced Industrial Science and Technology, AIST Tsukuba Center 6-13, Higashi 1-1-1, Tsukuba, Ibaraki 305-8566, Japan; Department of Medicine, Division of Medical Genetics, University of Washington, Seattle, Washington, 98195-7720, USA

**Keywords:** aging, homeostasis, age-related gene expression, blood coagulation, liver, FIX

## Abstract

HnRNP A3 is a protein that binds the age-related increase element (AIE) of blood coagulation factor IX (FIX) and that plays critical roles in age-related gene expression, likely through an epigenetic mechanism as yet unidentified. In a previous study, we found that Ser^359^ phosphorylated hnRNP A3 proteins do not bind to the AIE of FIX although both unphosphorylated and Ser^359^ phosphorylated hnRNP proteins exist in the liver. In the present study, to explore the relationship between hnRNP A3 and FIX, we examined the age-related expression pattern of 14 single spots of hnRNP A3 detected by 2DE and subsequent MALDI-TOF/TOF/MS analyses in mouse liver. We found that the level of all four Ser^359^ phosphorylated hnRNP A3 proteins increased with age (from 1-21 months), while the 10 unphosphorylated hnRNP A3 proteins showed various expression patterns with age. We then examined the functional role of hnRNP A3 in FIX expression using siRNA knockdown technology targeting the hnRNP A3 gene in aged mice (12-17 months old). Inhibition of hnRNP A3 expression induced an increase in the circulating FIX level in aged mice.

These results suggested that hnRNP A3 inhibits age-related FIX protein expression and that age-dependent modification of hnRNP A3, including its phosphorylation at Ser^359^, might be involved in the age-dependent increase in FIX expression *in vivo*.

## Introduction

The age-related stability element (ASE)/ age-related increase element (AIE)-mediated genetic mechanism for the regulation of age-related gene expression is the first molecular mechanism to occur in age-related homeostasis [1-3]. In this mechanism, two genetic elements, designated as ASE and AIE, play essential roles in the generation of age-related stable and age-related increases in gene expression, respectively. We recently reported that the Ets1 transcription factor and hnRNP A3 bind to ASE and AIE, respectively [4-5].

Regulation of AIE function is important because AIE functions in the regulation of age-related patterns of increased gene expression, which may contribute to the occurrence of age-related disease. The AIE was originally identified in the human blood coagulation factor IX gene (hFIX). The FIX gene has an 102 base pair (bp) stretch of dinucleotide repeats (rich in AT, GT and CA) in the middle of a 3′-untranslated region (3′-UTR), which potentially forms and functions as a stem-loop RNA structure (hereafter referred as hFIX-AIE RNA) after the gene is transcribed [1, 2, 6]. The mouse FIX gene (mFIX) also shows an age-related pattern of increased gene expression [1], and has a stretch of dinucleotide repeats (rich in AT) of about 50 bp in the middle of a 3′-UTR, potentially forming a stem-loop RNA structure (referred as mFIX-AIE RNA) after gene transcription [2]. Both hFIX-AIE RNA and mFIX-AIE RNA function equally well in a position-dependent and orientation-independent manner in producing an age-related increase pattern of gene expression, as tested with the human protein C gene, which lacks AIE [2].

The hnRNP A3 mRNA level in the mouse liver remains stable throughout the tested period of age (1 through 21 months) [5]. In contrast, the concentration of hnRNP A3 protein in the liver gradually increases in an age-dependent manner [5]

This discordance in the age-related expression patterns of the mRNA and protein of hnRNP A3 indicates that hnRNP A3 is regulated by age-dependent epigenetic events. Using 2DE and subsequent MALDI-TOF/TOF/MS analyses, our previous data showed 14 single spots for the hnRNP A3 protein at 3 months of age, with no contamination of other proteins, suggesting that hnRNP A3 is modified, including modification by phosphorylation at its Ser^359^. However, at present the detailed underlying mechanism of action of hnRNP A3 *in vivo* remains unknown.

In this study, to determine the age-related regulation of FIX by hnRNP A3 *in vivo*, the age-related profile of hnRNP A3 protein expression and the functional role of hnRNP A3 in the expression of FIX were investigated.

## Materials and Methods

### Animals

For the study of the effect of siRNA against hnRNP A3 on hFIX expression *in vivo*, animals carrying the minigene -802FIXm1/1.4 which has an additional 32 bp extension of the 5’ region of the hFIX gene that extends the region to nt -802 (containing the ASE region), and a 3’-UTR (containing the AIE region), were used [1]. The hFIX transgenic mice were constructed and subjected to longitudinal analyses as previously described [1, 4, 5]. C57BL/6J mice were used for 2DE and subsequent MALDI-TOF/TOF/MS (MALDI-TOF2) analyses of liver nuclear extracts (NEs). Animal care and use were reviewed and approved by the Committee for Animal Experimentation of the National Institute of Advanced Industrial Science and Technology (AIST)(permission # 36-07-009) and were performed in accordance with the institutional guidelines of the Committee for Animal Experimentation in the AIST.

### 2DE analysis of mouse liver nuclear proteins

Age-related expression analysis of hnRNP A3 proteins was carried out using an age-related mouse liver nuclear protein database that we previously made [7, 8, 9]. The image of 2DE gel electrophoresis that we previously reported [7] and that is shown in Fig. 1A was used for identification of the hnRNP A3 protein spots. Briefly, the 2DE gel electrophoresis analysis was performed as follows [7]. Mice (C57BL/6J mice) were maintained for at least one week on a 12 h light/12 h dark (LD) cycle with lights on from 6:00-18:00. Mouse liver was sampled during the time from 9:30 to 12:00. NEs (300 μg) were prepared from mouse liver tissues at 1, 3, 6, 12, 18 and 21 months of age and were subjected to IEF using a 24 cm immobiline DryStrip (pH 6-11) (BIO-RAD, Hercules, CA), followed by SDS-PAGE using pre-casted 10-16% gradient polyacrylamide gels (BIO-RAD) [1, 4, 5, 7, 8, 9]. The gels were then stained with Colloidal Coomassie Brilliant Blue (CBB) and scanned using a GS-800 scanner (BIO-RAD). Protein spots were analyzed using PDQuest software (BIO-RAD, version 7.1), and excised, destained in 50 mM ammonium bicarbonate/50% acetonitrile and dried in a vacuum concentrator (Savant, Holbook, NY) [5, 7]. Dried gel pieces were then rehydrated with 5 μl of 20 ng/L trypsin in 10 mM ammonium bicarbonate, and incubated at 30 °C overnight. The peptide containing fractions were collected and dried and were mixed with CHCA matrix before application to an Axima® CFR MALDI-TOF/MS (Shimazu Biotech, Kyoto, Japan) or an AXIMA® MALDI-TOF2 (Shimadzu Biotech). Peptide mass fingerprinting (PMF) analysis was performed using the MASCOT server version 2.1 (Matrix Science, London, U.K.) taking UniProt or NCBI databases as a reference database. Age-related expression analysis of the hnRNP A3 proteins of these spots in the present study was carried out using an age-related mouse liver nuclear protein database as previously reported [7, 8, 9].

**Figure 1.**
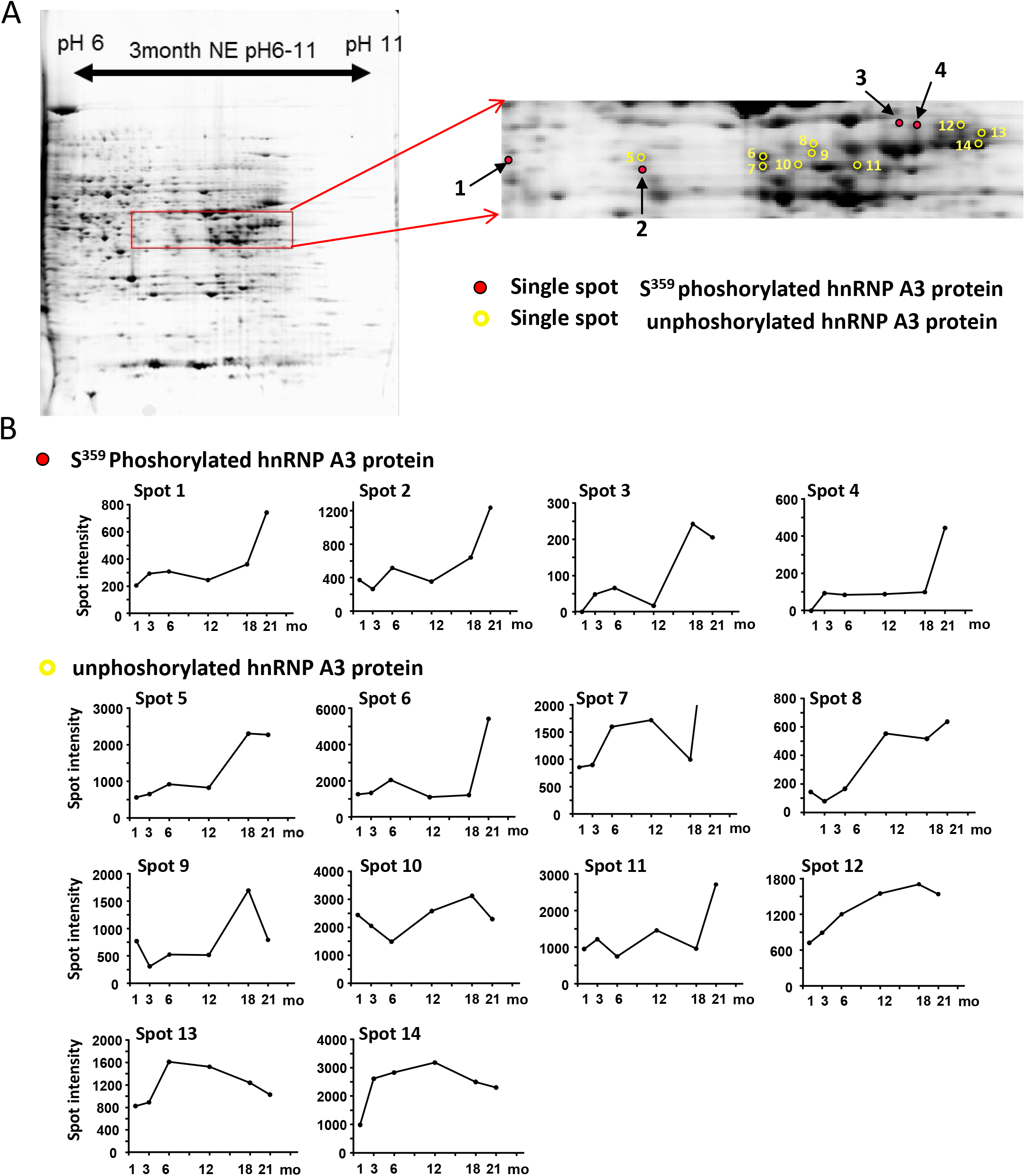
2DE analysis of hnRNP A3 proteins in the mouse liver with age. **A**. 2DE gel electrophoresis of mouse liver nuclear proteins at 3 months of age [7]. The spots of hnRNP A3 protein detected within the red square are magnified in the figure on the right. Single spots of Ser^359^ phosphorylated hnRNP A3 and unphoshorylated hnRNPA3 protein are marked in red and yellow circles respectively. **B**. Age-related expression profiles of each spot identified as hnRNP A3. Liver nuclear proteins of 1, 3, 6, 12, 18 and 21 month old mice were analyzed.

### Construction of a recombinant adenovirus harboring siRNA

An siRNA vector (siRNA 6) targeted against hnRNP A3 was prepared as previously reported [5]. Recombinant adenoviral particles harboring an siRNA expression unit against the target mRNA at a concentration of 8.4 ×10^9^ pfu in 100 μl PBS were administered into the tail veins of mice (hFIX transgenic mice carrying the minigene -802FIXm1/1.4) at 12 or 17 months of age.

#### Liver FIX mRNA Assay

Animals were sacrificed on day 4 or 14 post-viral injection and their liver tissues were removed, quick frozen on dry ice, and stored at –80 °C until use. Total RNA was prepared from these tissues using an acid guanidium phenol chloroform procedure. PCR primers and the TaqMan probe used for analysis of mFIX in these quantitative real-time RT-PCR assays were 5’-TGTCAGGAAAGCTGCAGGTTACT-3’ and 5’-CATCGTTCCTTCTGTGGCAGATT-3’, and 5’-FAM-CATCGTGCACCGCAA-MGB-3’, respectively. Similarly, for analysis of the β–actin (control), 5’-TCCACCTTCCAGCAGATGTG-3’ and 5’-CAGTAACAGTCCGCCTAGAAGCA-3’, and 5’-FAM-CATCGTGCACCGCAA-MGB-3’ were used, respectively. RT-PCR results were normalized to that of β–actin mRNA.

#### Assay of the circulating FIX level

The hFIX transgenic mice carrying the minigene -802FIXm1/1.4 were injected with recombinant adenoviral particles harboring the siRNA expression unit targeted against hnRNP A3 at 12 or 17 months of age. The FIX protein was then assayed as follows. Blood samples were individually collected (15 μl aliquots) via tail-tip snipping 2 days before and 3, 6, 9, 12 and 14 days after adenoviral particle injection. The serum obtained was routinely used to quantify circulating hFIX levels using a duplicated hFIX-specific ELISA as previously described [1, 4, 5].

## Results

### 2DE analysis of mouse liver nuclear extracts with age, and of spots originating from the hnRNP A3 protein

In previous studies, we identified 14 single spots of hnRNP A3 protein in liver NEs of mice at 3 months of age as shown in Fig. 1A [5, 7]. This multitude of spots is produced by post-translational modification of the hnRNP A3 protein and/or by alternative splicing. Our previous report showed that Ser^359^ of the hnRNP A3 protein is phosphorylated and that Ser^359^ is one of the sites of post-translational modification of hnRNP A3 in the mouse liver [5]. To elucidate other possible post-translational modifications of the hnRNP A3 protein in mouse liver, we studied the protein expression of the methylation enzyme, PRMT1, in mouse liver extracts with age, using western blot analysis (Supplemental Fig. S1). The protein level of PRMT1 was seen to increase along with age. PRMT1 is known to methylate the hnRNP proteins, A2/B1, A1, L [10, 11]. Western blotting indicated that the levels of these hnRNP proteins also increased with age although the hnRNP mRNAs, as assessed using gene chip analysis, were stably expressed with age (Supplementary Fig. S1 and S2). These results suggested the possibility that hnRNP A3 may also be methylated with age and that therefore methylation of hnRNP A3 may be involved in the age-related expression of FIX. Unfortunately, we did not obtain any evidence for modification of the hnRNP A3 protein by modification other than phosphorylation at Ser^359^ in the present MALDI-TOF/TOF/MS analyses, perhaps because our system might not be sensitive enough to detect a minute amount of modification or because the level of methylation is quite low.

In a previous study, we showed that the unphosphorylated hnRNP A3 protein binds to the AIE of FIX while the hnRNP A3 protein that is phosphorylated at Ser^359^ (which we refer to as Ser^359^ phosphorylated hnRNP A3) does not bind to AIE [5]. In the present study, to explore the age-dependent role of the hnRNP A3 protein in the mouse liver on FIX expression *in vivo*, we examined the hnRNP A3 protein profile with age using 2DE and subsequent MALDI-TOF/TOF/MS analyses, focusing on its Ser^359^ phosphorylation. In Figure 1A, the obtained spots of the hnRNP A3 protein are shown in the square window and are magnified. Four of the 14 hnRNP A3 single protein spots contained hnRNP A3 with phosphorylated Ser^359^, while the other spots contained hnRNP A3 with unphosphorylated Ser^359^ [5]. MS/MS analysis of the peptide fragment (aa 356 through 377, SSGSPYGGGYGSGGGGSGGYGSR) from the mass peak of m/z 1990.8 indicated that the Ser^359^ in the previously described peptide [5] is the post-translational phosphorylation modification site of hnRNP A3 in the livers of mice at 3, 6 and 21 months of age as shown in Supplemental Figure S3A, B, C.

In this study, we explored the liver NE content of the 14 single spots of the hnRNP A3 protein with age. We first examined the liver content of the four single spots of the Ser^359^ phosphorylated hnRNP A3 protein, which are marked in red in Fig. 1A (spots 1-4), with age (1-21 months). The Ser^359^ phosphorylated hnRNP A3 proteins in spot 1 and spot 2 were relatively highly expressed and showed an age-related increase in expression (Fig. 1B). The expression of the hnRNP A3 proteins in spot 3 and spot 4 also increased along with age although the expression of these proteins was low. Our previous report also indicated that the number of hnRNP A3 protein spots that were phosphorylated increased with age [7]. These previous data supported the results of the present study that the levels of hnRNP A3 proteins that are phosphorylated at Ser^359^ increase with age.

Unphosphorylated hnRNP A3 protein spots are marked in yellow in Fig. 1A. The Ser^359^ unphosphorylated hnRNP A3 proteins showed various expression patterns with age. Thus the expression of the hnRNP A3 proteins in spots 5, 6, 8, 9, 11 and 12 increased along with age (Fig. 1B), whereas the hnRNP A3 proteins in spot 13 and spot14 decreased with age. The expression of the hnRNP A3 proteins in spots 7 and 10 varied with age (Fig. 1B). Supplementary Figure 4 shows the expression of HuR proteins, which were used as a control, with age. Our previous report showed that HuR expression is stable with age as assessed by western blot analysis. The two spots of HuR that we detected in this study also showed stable expression with age.

### Effects of siRNAs against hnRNP A3 on FIX expression in the mouse liver *in vivo*

Next, we examined the functional role of hnRNP A3 in FIX expression using siRNA knockdown targeting the hnRNP A3 gene in aged mice (12-17 months old). FIX mRNA levels in the liver and FIX protein circulation levels in hFIX transgenic mice were examined. This study showed that animals that carried the hFIX minigene with both ASE and AIE showed increased FIX mRNA and protein expression with age (Fig. 2A). Our previous study showed that an siRNA vector against hnRNP A3 (siRNA6) reduced the hnRNP A3 mRNA level by approximately 78% in 293T cells *in vitro* and, furthermore, that delivery of adenoviral vectors harboring siRNA 6 into the tail veins of mice significantly reduced the hnRNP A3 mRNA level in the liver on day 4 and on day 14 *in vivo* [5]. In the present study, mouse liver transfected with adenoviral vectors harboring siRNA 6 against hnRNP A3 or its scrambled siRNA control at a concentration of 8.4 ×10^9^ pfu in 100 μl PBS, showed no reduction in hFIX mRNA expression over that of the cells treated with PBS-saline (control) on day 4 (Fig. 2B). However, on day 14, hFIX mRNA in the liver transfected with the adenoviral vector harboring siRNA 6 showed a considerable and significant increase compared to the level in the scrambled siRNA control. Next, we examined the hFIX protein level in the circulation after injection of the adenoviral vector harboring siRNA 6 over a period of 14 days (Fig. 2C, D, E, F). Animals transfected with adenoviral vector harboring siRNA 6 showed a gradual increase in circulating hFIX over time (Fig. 2C) with the highest increase observed on day 14. A scrambled siRNA or saline injection had no effect on the hFIX circulation level (Fig. 2D, E). This increase in the hFIX circulation level by injection with an adenoviral vector harboring siRNA 6 paralleled the change in hFIX mRNA levels in the liver after injection with an adenoviral vector harboring siRNA 6 (Fig. 2B and C). We therefore concluded that it was the hnRNP A3 itself that suppressed the expression of hFIX *in vivo*.

**Figure 2.**
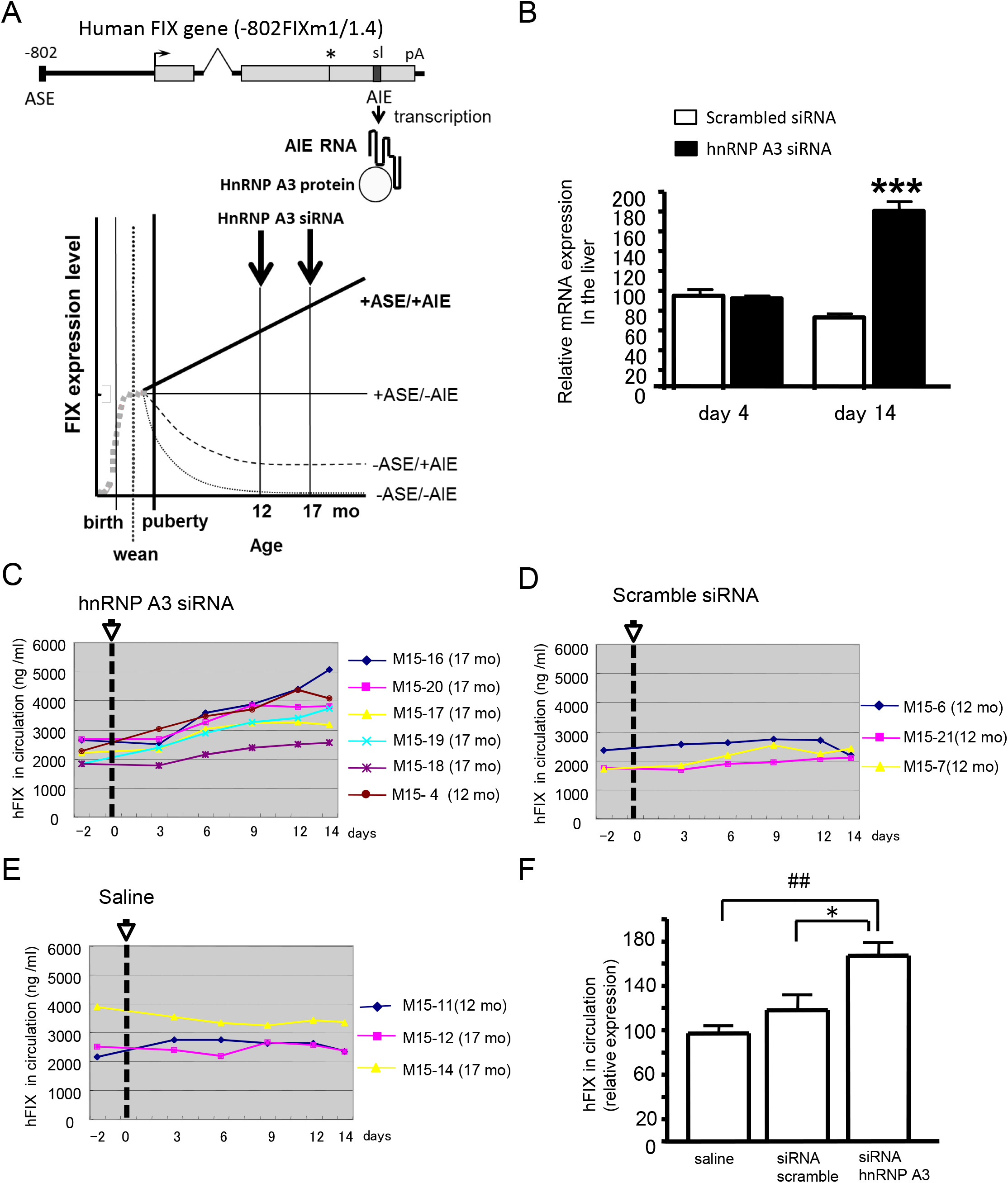
Effect of siRNAs against hnRNP A3 on FIX expression in the mouse liver *in vivo*. **A**. At the top, a schematic outline of the relationship between a key structural element (AIE) of the hFIX minigene (−802FIXm1/1.4) and the hnRNP A3 protein is shown. The promotor region (thick horizontal line at left), the 5’ terminal nt number and the ASE element are indicated. Transcribed hFIX regions are shown as gray rectangles. The shortened first intron is shown as a thin peaked line and the 3’ flanking sequence region is shown as a thick horizontal line at right. HnRNP A3 binds to the AIE located in the 3’ UTR of the FIX gene, which forms a stem-loop structure after gene transcription [1, 8, 9]. Right-pointing arrow, transcription start site; asterisk, translation stop site; pA, polyadenylation; sl, potential stem loop-forming dinucleotide repeats. At the bottom, four patterns of age-related hFIX minigene expression are shown. These previously reported four age-related mini-gene expressions were generated by various combinations of ASE and AIE, whose absence or presence is depicted with – or +, respectively [1, 8, 9]. Injection of the adenoviral vectors harboring siRNA against hnRNP A3 into mice carrying the hFIX minigene (−802FIXm1/1.4) are shown by vertical arrows at 12 or 17 months of age. Expression of the hFIX minigene was determined using ELSA [1, 4, 5]. **B**. Relative FIX mRNA levels in the mouse liver post-injection of adenoviral vectors harboring siRNA against hnRNP A3. The animals were injected with the adenoviral vectors harboring siRNA, scrambled siRNA, or PBS-saline, and the livers were sampled on day 4 and day 14 post-injection. Relative FIX mRNA expression levels in the liver, as determined using quantitative real time PCR are shown. The results were normalized to that of β-actin mRNA, and their mean relative values over that of the PBS-saline condition are presented. Thin vertical lines with short horizontal bars represent S.D. ***p <0.001 vs. Scrambled siRNA, Student’s *t*-test. **C. D. E**. FIX protein content in the plasma of individual mice before and after adenoviral vector injection. **C**, vector harboring siRNA against hnRNP A3. **D**, vector with scrambled siRNA. **E**, saline injection. **F**. Quantification of the effect of adenoviral siRNA against hnRNP A3 on the circulating FIX protein level on day 14 post-injection (## p<0.01, *p<0.05, Dunnet’s test).

## Discussion

HnRNP A3 is known to be involved in mRNA decay and shuttling [12]. Dysregulation of the functions of hnRNP A3 have been linked with multiple diseases [13-15]. Our previous analysis of mouse liver nuclear proteins by 2DE and subsequent MALDI-TOF/TOF/MS indicated that the hnRNP A3 protein has many post-translational modifications [5, 7]. There are many supporting papers for hnRNP A3 protein modification by methylation, phosphorylation and sumoylation [10, 11, 16, 17]. Among the age-related post-translational modifications of the hnRNP A3 protein is phosphorylation of the hnRNP A3 protein at Ser^359^ in the mouse liver. The phosphorylation of hnRNP proteins is important for activation of the hnRNP proteins themselves [18-23]. Our present analysis of these 2DE spots indicated that the level of Ser^359^ phosphorylated hnRNP A3 proteins increased along with age and correlated with the age-related increase in expression of FIX [7]. In this respect it is interesting that hnRNP A3 proteins that are not phosphorylated at Ser^359^ bind to the AIE of FIX, while Ser^359^ phosphorylated hnRNP A3 proteins do not bind to the AIE. Our previous study showed that the number of spots that originated from the Ser^359^ phosphorylated hnRNP A3 protein increased in old age (18 and 21 months) in mice [7]. The phosphorylation and dephosphorylation of this region of the hnRNP A3 protein is likely to play critical roles in the regulation of FIX mRNA translation and in the intracellular recycling of hnRNP A3 itself. Further detailed studies of the age-related regulation of the FIX gene by phosphorylation and dephosphorylation of hnRNP A3 and of the underlying mechanism of action of hnRNP A3 remain to be carried out.

The ASE/AIE-mediated genetic mechanism for age-related gene regulation has two essential elements, ASE and AIE [1], which bind to Ets1 and hnRNP A3, respectively [4-5]. In this study, to evaluate the effect of knockdown of hnRNP A3 expression in aged mice using the highly effective siRNA, siRNA 6, which we reported previously [5], we examined the FIX mRNA content in the liver and the circulating level of FIX in aged mice after administration of siRNA 6 *in vivo*. Our previous report showed that this siRNA lowered the hnRNP A3 mRNA level *in vitro* to 49% and 54% of the level in PBS or scrambled siRNA controls on day 4 and day 14 after infection, respectively [5]. However, in the present *in vivo* study FIX mRNA expression was significantly increased only on day 14 after siRNA 6 administration FIX protein in the circulation, which originates in the liver, also gradually increased after siRNA 6 administration. This effect of siRNA 6 was specific, because no such effect was seen with saline or scrambled siRNA (Fig. 2D, E). Interestingly, these results are in contrast to our previous in vitro study in which we demonstrated that the hnRNP A3 protein binds to the AIE of the FIX gene and that hFIX expression in HepG2 cells carrying -416FIXm1/1.4 showed siRNA6-specific and adenoviral MOI-dependent reductions in hFIX expression [5].

We could not think of any reason why FIX expression was increased on day 14 following siRNA 6 injection in our present *in vivo* study. One possible explanation is that the hnRNP A3 protein binds to FIX mRNA and only allows initiation of translation after hnRNP A3 is phosphorylated at Ser^359^(Fig.3). Some hnRNP proteins that bind to mRNA have been reported to allow initiation of translation after their phosphorylation [19, 20, 23]. For example, hnRNP K, which is an hnRNP family protein, is known to bind to the differentiation control element (DICE), a CU-rich stretch of sequence in the 3’-UTR of 15-lipoxygenase (LOX) mRNA, thereby inducing translational repression of the LOX gene [18]. This repressive effect is removed upon tyrosine phosphorylation of the hnRNP K [19]. Zipcode binding protein 1 (ZBP1), which binds to a conserved 54-nucleotide region in the 3’-UTR of β-actin mRNA, prevents the mRNA from being translated, but again, like hnRNP K, phosphorylation of Tyr^396^ of hnRNP K frees the bound mRNA, allowing it to be translated [20]. In our previous study, hnRNP A3 that was bound to the AIE RNA in the FIX mRNA was not phosphorylated at Ser^359^, while hnRNP A3 in the mouse liver nuclear extracts was a mixture of Ser^359^phosphorylated and unphosphorylated forms [5]. *In vivo*, an age dependent increase in phosphorylated hnRNP A3 occurs in the mouse liver, and thus FIX mRNA expression may be regulated by phosphorylated/non-phosphorylated hnRNP A3 through a similar mechanism by which HnRNP K or ZBP1 regulate the mRNAs to which they bind.

**Figure 3.**
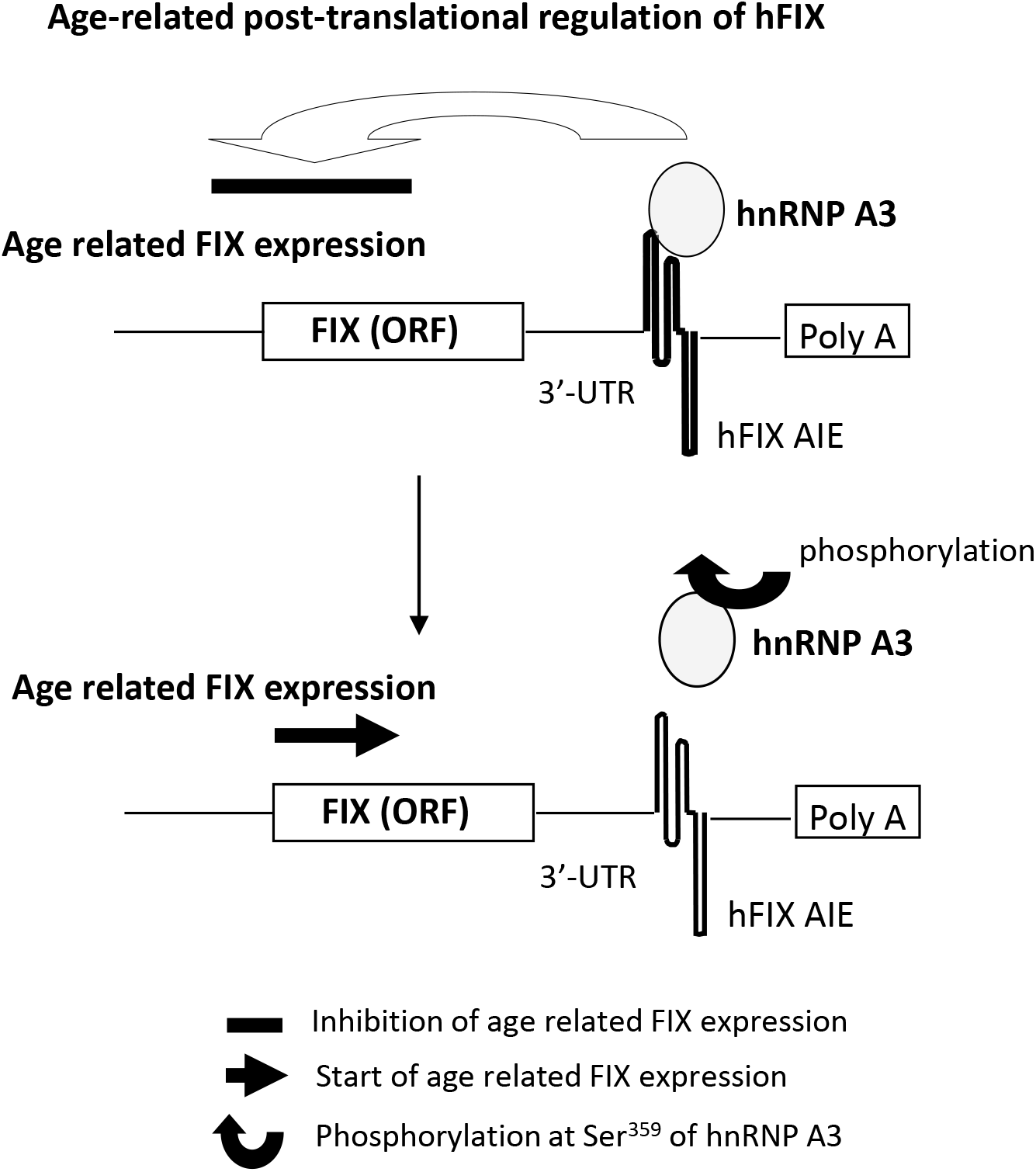
Hypothesis of the age-related regulation of FIX expression by the hnRNP A3 protein. Hypothesis of the involvement of the hnRNP A3 protein in the increase in FIX expression with age is shown. (Top) Binding of the hnRNP A3 protein to the AIE of FIX mRNA inhibits the age-related increase in FIX expression in an age dependent manner. (Bottom) This repressive effect of hnRNP A3 is removed upon the phosphorylation of hnRNP A3 at Ser^359^ by a mechanism similar to that which has been reported in studies of hnRNP K, A1 and ZBP1 [15. 16. 19].

The results of the present study clearly showed that the hnRNP A3 protein has an inhibitory effect on FIX protein expression and that age-dependent phosphorylation of hnRNP A3 at Ser^359^ might be involved in the age-dependent increase in FIX expression *in vivo*.

## Acknowledgments

This research was in part supported by the internal research fund of AIST and was partially supported by JSPS KAKENHI Grant Number 24621001. We thank Dr. Kenneth Sutherland for comments on drafts of this manuscript.

## Author Contributions

Conceived and designed the experiments: TH. Performed the experiments: TH HK SK. Analyzed data: TH HK SF SK. Contributed reagents/materials/analysis tools: TH SK. Wrote the paper: TH.

## Figure Legends

**Supplementary Fig. S1.**
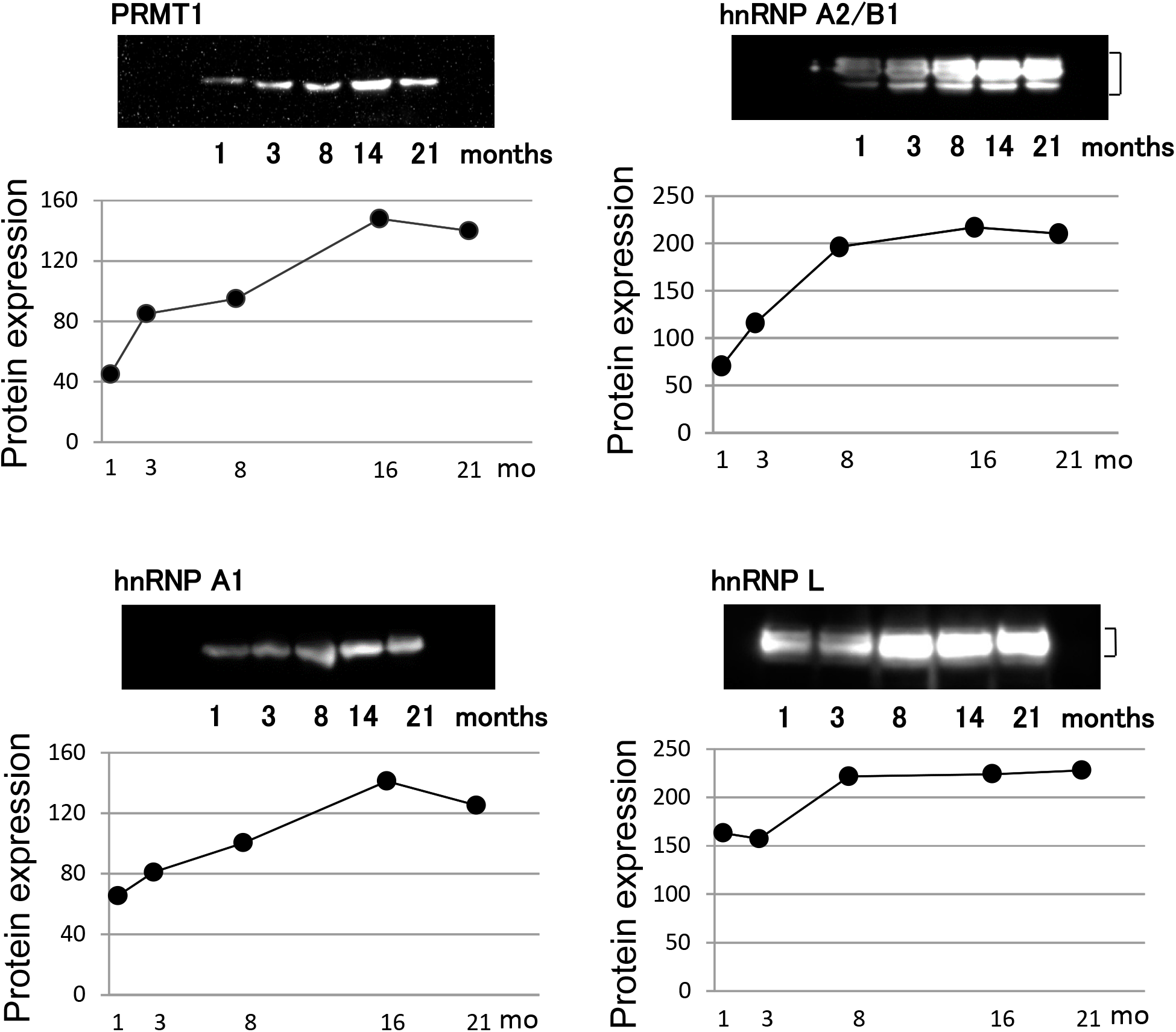
Age-related expression of PRMT1, hnRNPA2/B1, hnRNPA1 and hnRNPL proteins in mouse liver. Western blot analysis of the protein expression of PRMT1 and hnRNP family proteins in liver nuclear extracts from C57BL/6 mice aged 1, 3, 8, 16, and 21 months (mo) is shown. Samples were prepared as previously described [4, 5, 7]. Proteins were solubilized in 20 μL of SDS loading buffer. After adjustment of the protein concentration using the BCA method, the samples were subjected to 10% SDS-PAGE. Western blot analyses were carried out according to standard methods, using a horseradish peroxidase–conjugated anti-rabbit secondary antibody for signal detection with the ECL assay system (GE Healthcare). Band intensity was measured using a Fujix Bio-imaging analyzer LAS 1000 Ver. 3.4X software (Fujifilm, Tokyo, Japan). Mouse anti-hnRNP A2/B1 (mouse monoclonal; DP3B3, ImmuQuest, Cleveland, UK), anti-hnRNP A1 (mouse monoclonal; 4B10, ImmuQuest), anti-hnRNP L (mouse monoclonal; 4D11, ImmuQuest) and rabbit polyclonal anti-PRMT1 (upstate, Lake Placid, NY) antibodies were used as the first antibodies. Protein expression of PRMT1 and hnRNP family proteins is shown as the mean intensity of each band at each age.

**Supplementary Fig. S2.**
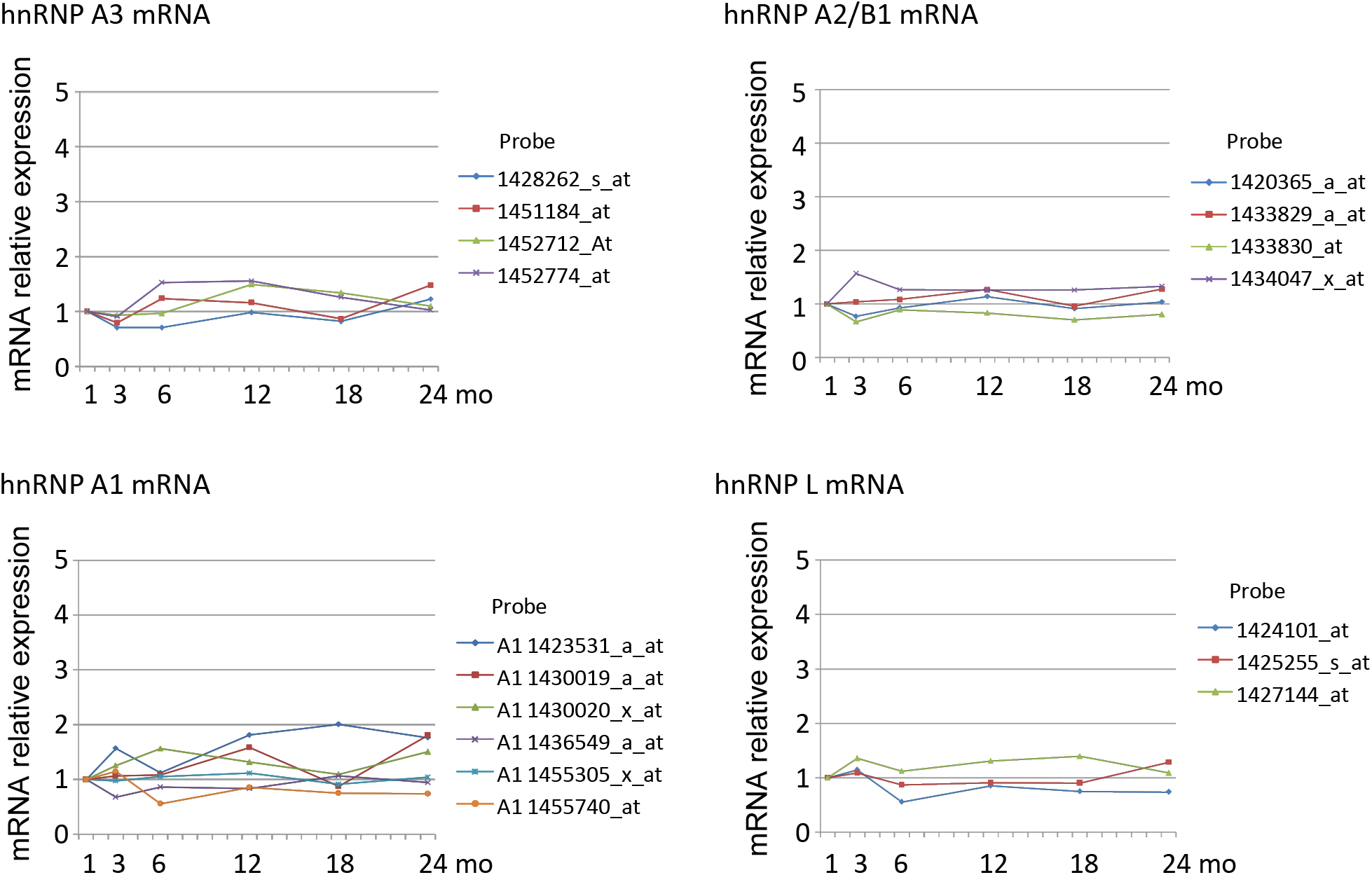
Age-related expression of hnRNP A3, hnRNP A2/B1, hnRNP A1 and hnRNP L mRNAs in mouse liver. The mRNA expression of hnRNP A3, hnRNP A2/B1, hnRNP A1 and hnRNP L in the liver of mice at different ages was measured using gene chip analysis. The GeneChip® Murine Expression Array 430A (MOE430A) (Affymetrix, Santa Clara, CA) was used for profiling the expression of the indicated mouse liver genes at various ages. Total RNA from liver lobes of mice aged 1, 3, 6, 12, 18 and 24 months was isolated using the TRIzol® Reagent (Invitrogen, Carlsbad, CA). RNA preparations were processed for 30 min at 37 °C with RQ1 RNase-Free DNase (Promega, Madison, WI) according to the manufacturer’s instructions, and their quality was evaluated using the RNA 6000 Nano assay kit and Agilent 2100 Bioanalyser (Agilent Technologies, Palo Alto, CA). An aliquot of the total RNA (7 μg) was annealed to 100 pmol of a T7-oligo (dT) primer in a final volume of 12 μl, and was denatured by incubation at 70 °C for 10 min followed by cooling on ice for 2 min. The first strand was synthesized for 1 h at 42 °C with 200 units of SuperScript Reverse Transcriptase (Invitrogen) according to the manufacturer’s instructions. Second strand synthesis was then done for 2 h at 16 °C by using RNase H, E. coli DNA polymerase I, E. coli DNA ligase and a dNTP mix. The double stranded cDNAs obtained were then incubated with T4 DNA polymerase (19 units) for 5 min at 16 °C, followed by a clean-up process using 10 μl of 0.5 M EDTA of the GeneChip Sample Cleanup Module (QIAGEN, Valencia, CA). The resulting cDNA preparation was then used for *in vitro* transcription for 5 h at 37 °C using the Bioarray High Yield RNA Transcript Labeling Kit (Enzo Life Sciences, Farmingdale, NY). Biotinylated cRNAs were cleaned again using the GeneChip Sample Cleanup Module. The cRNA were then fragmented by incubation at 94 °C for 35 min in the fragmentation buffer, and the final fragmented cRNAs (15 μg) were hybridized to the chip MOE439A (Affymetrix) for 16 h at 45 °C as described in the manufacturer’s instructions. Stained arrays were scanned with a GeneArary scanner (Affymetris). Scanned array data were analyzed using the GeneSpring GX 7.3.1 program (Silicon Genetics, Redwood City, CA). GeneChip data at six age time points (1,, 3, 6, 12, 18 and 24 months) were loaded onto the GeneSpring software, and normalized per chip to the 50th percentile and per gene to 1 month samples.

**Supplementary Fig. S3.**
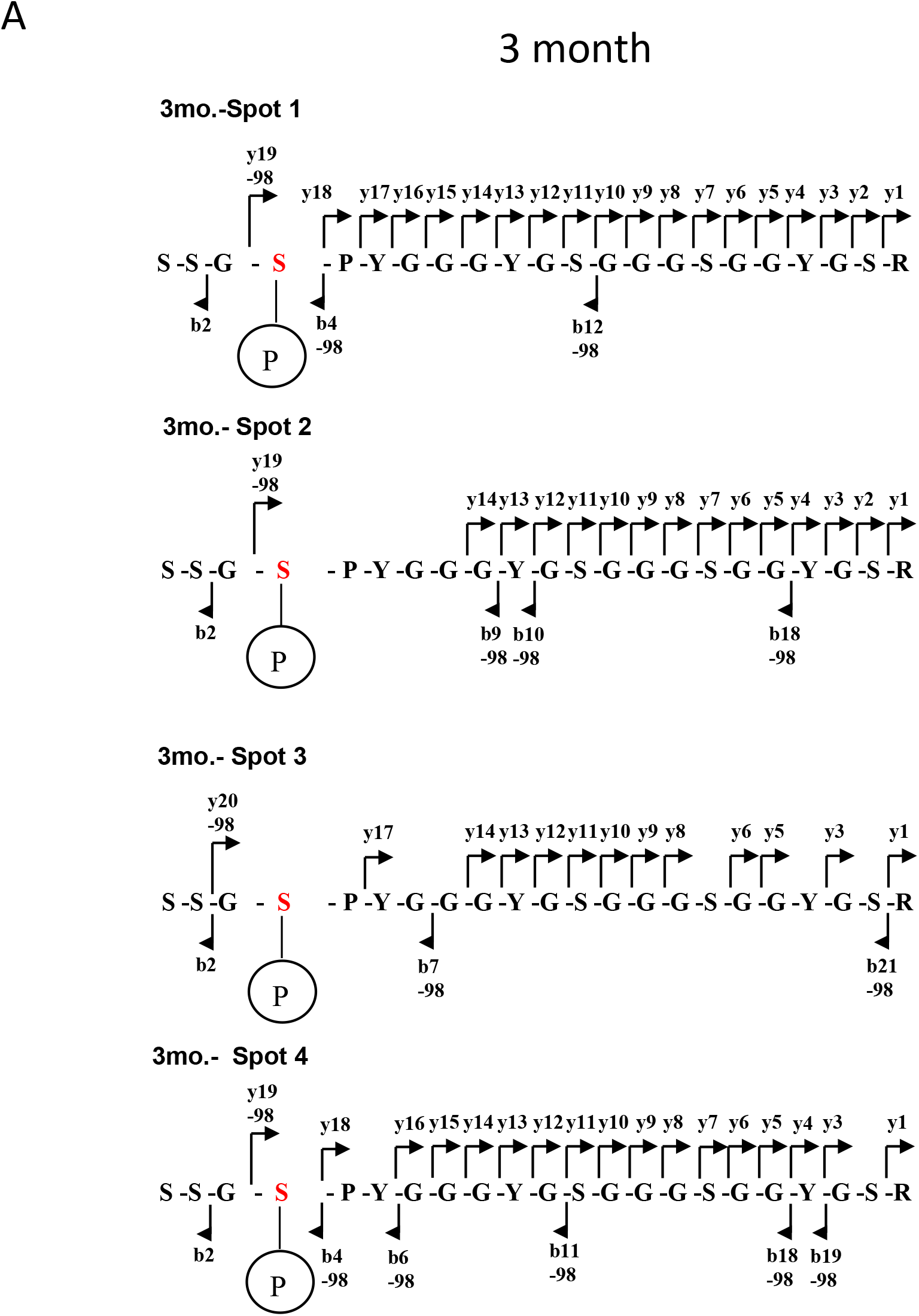
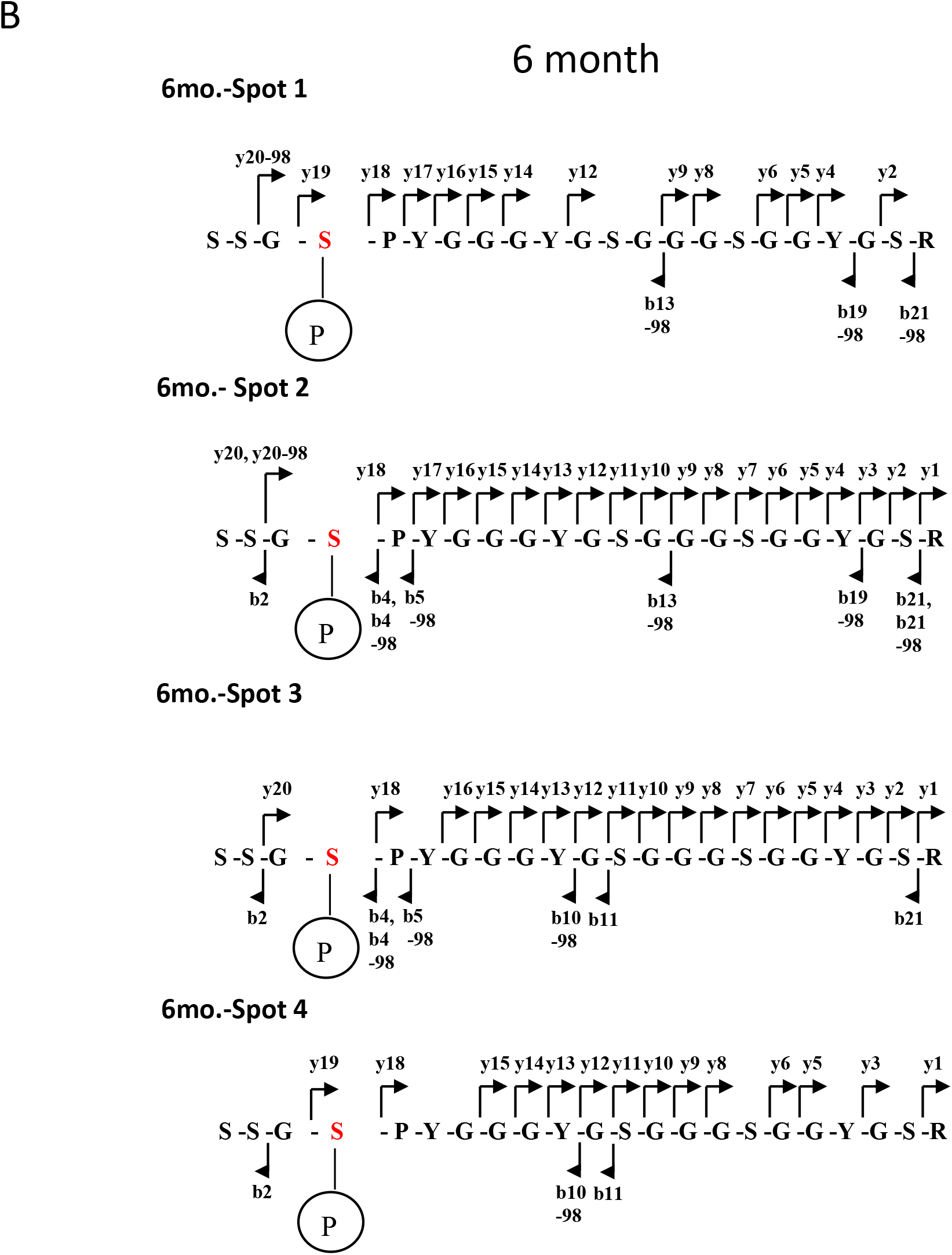
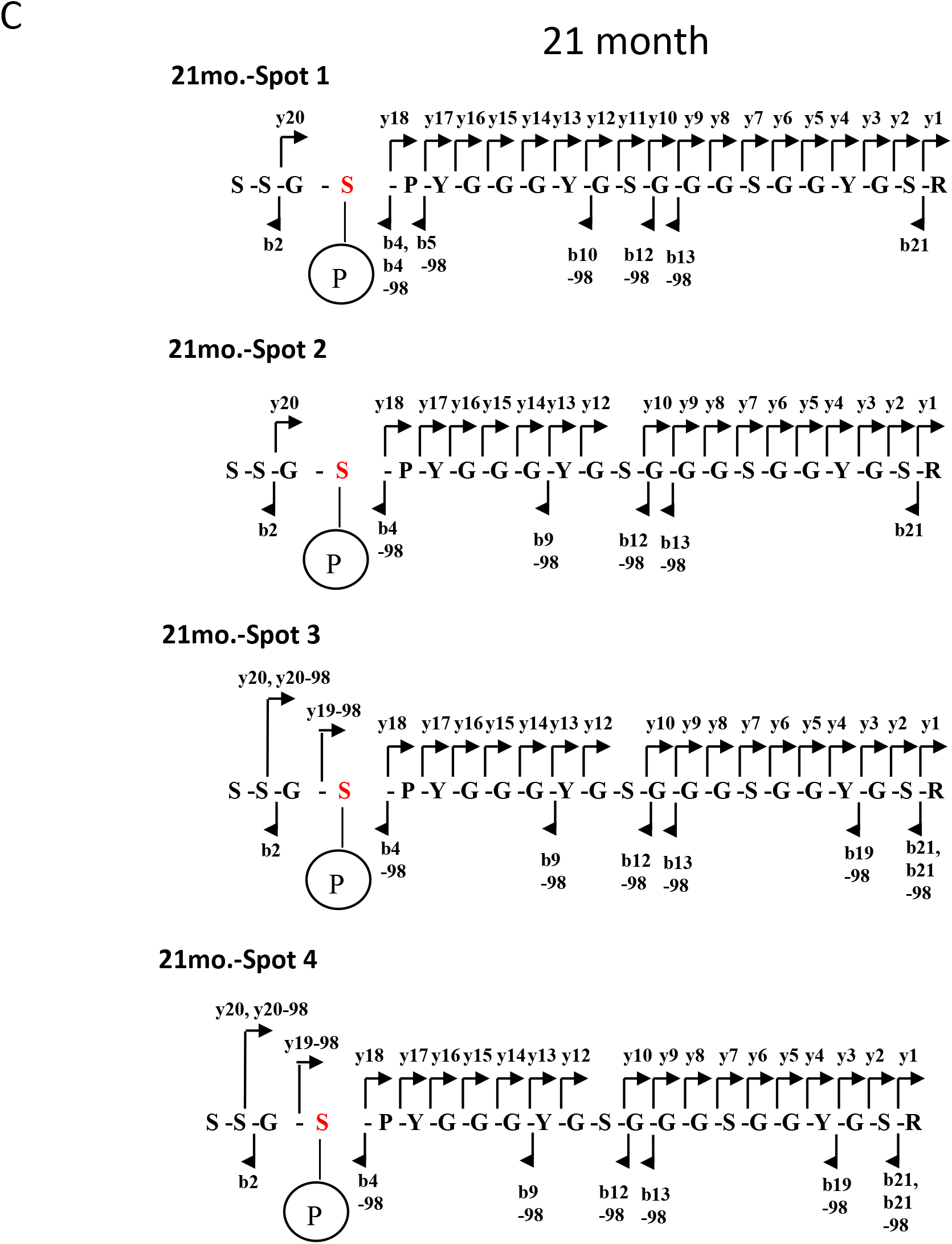
The phosphorylation status of the hnRNP A3 protein in liver nuclear extracts of mice aged 3, 6 and 21 months. Results of mass spectroscopy of hnRNP A3 spots. Annotated ions as nominal m/z values and the result of fragmentation of an amide bond with charge retention on the carboxyl end (y series ion) or amino terminus (b series ion) are shown. The fragment ions, y19-98, y20-98, b4-98, b5-98, b6-98, b7-98, b9-98, b10-98, b11-98, b12-98, b13-98, b18-98, b19-98 and b21-98 showed a loss of 98-Da in mass corresponding to a phosphate group. The fragment ions b2 and y18 indicated that a phosphate group is not present in the SS dipeptide sequence (Ser ^356^ through Ser ^357^) or in the sequence PYGGGYGSGGGGSGGYGSR (Pro ^360^ through Arg ^377^). These data indicated that Ser ^359^ (Red) is the only site in the peptide fragment SSGSPYGGGYGSGGGGSGGYGSR (aa 356 through 377) that is phosphorylated with age at 3, 6 and 21 months (A, B and C, respectively).

**Supplementary Fig. S4.**
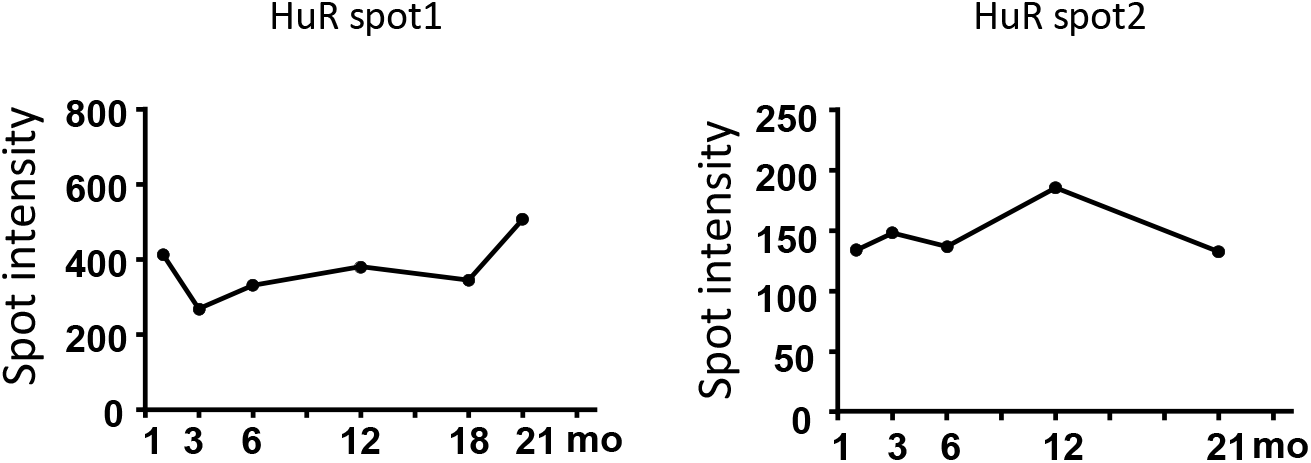
The expression of HuR proteins in mouse liver with age. Age-related expression profiles of spots identified as HuR protein are shown. Proteins of liver nuclear extracts of mice aged 1, 3, 6, 12, 18 and 21 months were analyzed. Two spots of HuR protein were detected in the present study.

